# Genome-wide SNP discovery in field and laboratory colonies of Australian *Plutella* species

**DOI:** 10.1101/141606

**Authors:** K.D. Perry, S.M. Pederson, S.W. Baxter

**Affiliations:** SCHOOL OF AGRICULTURE, FOOD & WINE, UNIVERSITY OF ADELAIDE, SOUTH AUSTRALIA 5005; SCHOOL OF BIOLOGICAL SCIENCES, UNIVERSITY OF ADELAIDE, SOUTH AUSTRALIA 5005

**Keywords:** Gene flow, population structure, RAD-Seq, *Plutella australiana*, insecticide resistance

## Abstract

Understanding dispersal and gene flow is an important focus of evolutionary biology, conservation biology and pest management. The diamondback moth, *Plutella xylostella*, is a worldwide pest of *Brassica* vegetable and oilseed cropping systems. This insect has high dispersal ability, which has important consequences for population dynamics and the potential spread of insecticide resistance genes. Population genetic studies of the diamondback moth have found little evidence of population structure, suggesting that frequent intermixing occurs within regions, however the patterns of local and regional dispersal remain to be identified. For this and many other pest species, understanding dispersal is crucial for developing integrated management tactics such as forecasting systems and insecticide resistance management plans. In recent years, next generation sequencing (NGS) methods have provided previously unparalleled resolution for population genetic studies in a wide range of species. Here, we assessed the potential of NGS-derived molecular markers to provide new insights about population structure in the diamondback moth. We use restriction-site-associated DNA sequencing (RAD-Seq) to discover hundreds to thousands of single nucleotide polymorphism (SNP) markers in nine field and laboratory-reared populations collected from Australia. Genotypic data from RAD-Seq markers identified a cryptic species, *P. australiana*, among individuals collected from a wild host, *Diplotaxis* sp., indicating strong divergence in the nuclear genomes of two Australian *Plutella* lineages. Significant genetic differentiation was detected among populations of *P. xylostella* used in our study, however this could be explained by reduced heterozogosity and genetic drift in laboratory-reared populations founded by relatively few individuals. This study demonstrates that RAD-Seq is a powerful method for generating SNP markers for population genetic studies in this species.

## INTRODUCTION

Dispersal is a fundamental life history trait with important consequences for the spatial and temporal dynamics of populations. ‘Effective’ dispersal (resulting in reproduction) also affects allele frequencies and the genetic structure of populations, and consequently, evolutionary processes (Broquet and Petit 2009). For example, adaptation and speciation depend on a balance between selection and gene flow (Turelli et al. 2001, Endersby et al. 2008). For pest species, quantifying dispersal and its effects on genetic structure is crucial to developing integrated management tactics, such as forecasting systems (Lushai and Loxdale 2004, Zalucki and Furlong 2005) and insecticide resistance management strategies. Molecular markers are a powerful tool for assessing the geographic structure of populations and inferring patterns of gene flow at large scales (Roderick 1996). Population genetic approaches have been successfully employed to infer patterns of long distance dispersal in a range of insect pests (Kim and Sappington 2013, Sun et al. 2015).

The diamondback moth, *Plutella xylostella*, is a worldwide pest of *Brassica* vegetable and oilseed crops. (Furlong et al. 2008, Zalucki et al. 2012, Furlong et al. 2013). The success of this insect is due in part to its remarkable genetic plasticity (Henniges-Janssen et al. 2011), large effective population sizes and high genetic diversity (You et al. 2013) that enable it to rapidly adapt to local environments and evolve insecticide resistance (Furlong et al. 2013). Furthermore, the diamondback moth is highly mobile and well-adapted to exploit its short-lived Brassicaceous hosts. It displays predominantly short-range dispersal within high quality host patches (Mo et al. 2003) but when habitat quality deteriorates, may utilize high altitude air currents to migrate long distances and colonise new habitats (Chu 1986, Chapman et al. 2002, Leskinen et al. 2011, Fu et al. 2014). In some years, large scale migration events instigate damaging outbreaks in crops (Dosdall et al. 2004, Wei et al. 2013). In most regions, the dispersal ecology of diamondback moth is poorly understood, yet this knowledge is critical for the development of forecasting systems and insecticide resistance management plans (Furlong et. al. 2008). One reason for this is the challenging nature of studying long distance dispersal in small insects (Lushai and Loxdale 2004).

Previous population genetic studies in diamondback moth have employed a range of molecular markers, including allozymes (Caprio and Tabashnik 1992, Noran and Tang 1996, Kim et al. 1999, Pichon et al. 2006), ISSRs (Roux et al. 2007), microsatellites (Endersby et al. 2006) and mitochondrial genes (Chang et al. 1997, Kim et al. 2003, Li et al. 2006, Saw et al. 2006, Niu et al. 2014). Several authors have reported population differentiation at inter-continental scales (Endersby et al. 2006, Pichon et al. 2006, Roux et al. 2007). However, most studies from around the world have found little evidence of population structure within regions, including China (Kim et al. 1999, Li et al. 2006), Korea (Kim et al. 1999, Kim et al. 2000, Kim et al. 2003, Li et al. 2006), the USA (Caprio and Tabashnik 1992, Chang et al. 1997) and Australia (Endersby et al. 2006, Saw et al. 2006). These findings suggests that frequent intermixing occurs within regions, however the local and regional patterns of dispersal remain to be identified. More recently, studies using mitochondrial markers, or complementing these with microsatellite (Wei et al. 2013) or ISSR (Yang et al. 2015) nuclear markers, have provided new insights into seasonal migration routes from southern to northern regions of China, and identified potential geographic barriers to gene flow (Niu et al. 2014). Mitochondrial markers have also recently identified a novel *Plutella* lineage in Australia (Landry and Hebert 2013).

In recent years, next generation sequencing (NGS) has revolutionized the fields of molecular ecology and population genetics. Reduced representation sequencing methods (Narum et al. 2013) combined with the power of high-throughput NGS platforms (Glenn 2011) facilitate rapid and cost-effective marker discovery and genotyping in a wide range of organisms (Davey et al. 2011). Restriction-site-associated DNA sequencing (RAD-Seq) (Baird et al. 2008) is one of several reduced-representation methods for sequencing targeted regions across the genome at high sequencing depth, providing numerous advantages over traditional markers for population genetic studies (Davey and Blaxter 2010). RAD-Seq studies have provided new insights into previously undetected population genetic structure in a wide range of contexts (Narum et al. 2013, Reitzel et al. 2013) including terrestrial invertebrates (Nadeau et al. 2013, Lozier 2014).

Here, we assess the potential of NGS methods to provide new insights about population structure in the diamondback moth. We use RAD-Seq to discover single nucleotide polymorphism (SNP) markers in nine field and laboratory-reared populations collected from Australia. The RAD-Seq markers facilitate an initial assessment of genetic diversity within and among these populations.

## MATERIALS AND METHODS

### Sample collection

Samples of diamondback moth were collected from *Brassica* vegetables, canola or wild *Brassica* hosts from nine locations in Australia between September 2012 and April 2014 (Table 1, Figure 1B). At each location, individuals were collected using a sweep net or by direct sampling. Seven populations were reared in laboratory cages on cultivated cabbage and 10% honey solution for between one and six generations. Individuals from two field populations and six of the laboratory-reared populations were preserved in 20% DMSO, 0.25M EDTA salt saturated solution (Yoder et al. 2006) and stored at −80°C. Individuals from the Nundroo population were stored in USP Grade propylene glycol and stored at −20°C.

### RAD library preparation and sequencing

Libraries for RAD sequencing were prepared following a protocol modified from Baird et al. (2008). Genomic DNA was extracted from individual larvae or pupae by homogenizing tissue in DNA isolation buffer (Zraket et al. 1990) followed by two phenol and one chloroform extractions. DNA was treated with RNase A then precipitated and re-suspended in TE buffer. Genomic DNA was quantified using a Qubit 2.0 fluorometer (Invitrogen) and 200 ng digested with 10 units of *Sbf1* in Cutsmart Buffer (NEB) for 1 hour at 37°C then heat inactivated at 80°C for 20 minutes. P1 adapters with one of six molecular identifiers (MIDs) (AATTT, AGCTA, CCGGT, GGAAG, GTCAA or TTCCG) were annealed then ligated to digested DNA (top strand 5’-GTTCAGAGTTCTACAGTCCGACGATCxxxxxTGCA-3’, bottom strand 5’-Phos-xxxxxGATCGTCGGACTGTAGAAC-3’, x represents sites for MIDs) using 1 μL T4 DNA ligase (Promega), 1 mM ATP, Cutsmart Buffer. Six individuals from different populations were pooled to form 12 library groups, each containing different P1 adapters to facilitate sample multiplexing. Library pools were then sheared using a Bioruptor sonicator (diagenode), ends blunted (NEB), adenine overhangs added then P2 adapters ligated (top strand 5’-Phos-TGGAATTCTCGGGTGCCAA-3’, bottom strand 5’-CCTTGGCACCCGAGAATTCCAT-3 ‘). DNA purification between each step was performed with magnetic beads (AMPure). PCR library amplification conditions were 16 cycles of 98°C for 10 seconds, 65°C for 30 seconds and 72°C for 30 seconds using RP1 (forward) 5’-AATGATACGGCGACCACCGAGATCTACACGTTCAGAGTTCTACAGTCCGA-3 ‘ and 12 unique RPI-indexed (reverse) 5’-CAAGCAGAAGACGGCATACGAGATxxxxxxGTGA CTGGAGTTCCTTGGCACCCGAGAATTCCA-3’ primers. Libraries were run on agarose gel to size select DNA fragments 300-700 base pairs in length. Paired end sequencing using 100 bp reads was performed over two lanes of Illumina HiSeq2500 at the Australian Cancer Research Foundation (ACRF) Cancer Genomics Facility.

### Read filtering and variant calling

A total of 131.6 million raw sequence reads were demultiplexed using radtools v1.2.4 (Baxter et al. 2011) then 50.3 million PCR duplicates were removed using the clone_filter tool in stacks v1.19 (Catchen et al. 2013). Read trimming, adapter removal and quality filtering were performed in trimmomatic v0.32 (Bolger et al. 2014). First, a thymine base overhang added during P2 adapter ligation was trimmed from reverse reads, then paired end trimming was performed using the ILLUMINACLIP tool to remove adapter, trailing low quality bases (quality score<3), bases within a 4-base sliding window with average quality below 15 and trimmed reads shorter than 40 bp. Paired reads were aligned to the *Plutella xylostella* reference genome (version 1.1, modified to include the mitochondrial genome, accession number: JF911819) using stampy v1.0.21 (Lunter and Goodson 2011) with --baq and -- gatkcigarworkaround options and expected substitution rate set to 0.005. Genotypes were called using the genome analysis toolkit (gatk) v3.3-0 (McKenna et al. 2010, DePristo et al. 2011) HaplotypeCaller tool following the GATK Best Practices Workflow for GVCF-based Cohort Analysis (Van der Auwera et al. 2013). Sites with a genotype quality (GQ) > 30 were retained. Filtering was performed using VCFTOOLS v0.1.12a (Danecek et al. 2011) to identify a set of variant sites for population genetic analysis. We removed indels and retained bi-allelic SNPs that passed the following quality filters: genotyped in at least 60 of 72 individuals, QUAL≥400, average read depth between 20 and 100 across individuals, minor allele frequency ≥ 0.2 and in Hardy-Weinberg equilibrium with p-value set to 0.05. To avoid closely linked markers, variants were separated by a minimum distance of 2 kb using the vcftools --thin function. A final set of 1285 SNP variants were retained after filtering. In addition, from the GATK HaplotypeCaller output, we generated a set of all confidently called variant and invariant sites (GQ≥30). Filtering was performed using VCFTOOLS v0.1.12a (Danecek et al. 2011) to remove indels and sites located within transposons, and retain sites genotyped in 60 of 72 individuals with mean depth between 20 and 100 across individuals. After filtering, we retained 491 831 confidently called variant and reference sites, including 623 sites from the mitochondrial genome.

**Table 1:** Collection details of *Plutella* populations from Australia genotyped using RAD-Seq

| Location | Latitude | Longitude | Host | Collection date | <sup>a</sup> Lifestage collected | <sup>b</sup> F<sub>1</sub> Individuals | Generation sequenced |
| --- | --- | --- | --- | --- | --- | --- | --- |
| Esperance, Western Australia | -33.8588 | 121.8931 | Canola | 26/9/2012 | L, P | 59 | F6 |
| Nundroo, South Australia | -31.7516 | 132.0565 | Canola | 19/8/2013 | L | 86 | F2 |
| Calca, South Australia | -33.0492 | 134.3729 | Wild *Brassica*<sup>c</sup> | 8/4/2014 | L | - | Field |
| Picnic Beach, South Australia | -34.1696 | 135.2744 | Wild *Brassica*<sup>d</sup> | 7/4/2014 | L | 40 | F1 |
| Mallala, South Australia | -34.4383 | 138.5099 | Canola | 11/9/2013 | L | 173 | F5 |
| Virginia, South Australia | -34.715 | 138.557 | Cauliflower | 7/3/2013 | L, P | - | Field |
| Glenore Grove, Queensland | -27.528 | 152.407 | Cabbage | 11/10/2012 | L, P | 25 | F5 |
| Mt Sylvia, Queensland | -27.717 | 152.220 | Cabbage | 3/10/2012 | L, P | 59 | F5 |
| Tenthill, Queensland | -27.566 | 152.235 | Red cabbage | 11/10/2012 | L, P | 40 | F5 |

<sup>a</sup> Larvae (L), pupae (P), <sup>b</sup> Number of F<sub>1</sub> individuals used to establish laboratory populations, <sup>c</sup> *Diplotaxis* sp., <sup>d</sup> Sea rocket, *Cakile maritima*

### Population genetic analysis

We used the 491 832 sites to generate a phylogeny for 72 individuals using a neighbor-joining clustering method implemented in the program geneious v7.1.9 (Kearse et al. 2012). An individual representing a cryptic species, *P. australiana*, ‘Calca-6’, was used as the out-group. We assumed the Tamura-Nei (1993) model and resampled with 1000 bootstraps to generate a consensus tree displaying nodes with at least 50% consensus support and visualized the resulting tree in the program figtree v1.4.2 (http://tree.bio.ed.ac.uk/software/figtree/).

Summary population statistics were calculated for variant SNPs (n=1285) and separately for all confident variant and invariant sites (n=491 832). The vcftools (Danecek et al. 2011) --depth function was used to calculate average site depth, and vcf-stats used to calculate the average number of genotyped sites and private alleles. Expected and observed heterozygosity and the inbreeding coefficient, F_IS_, were calculated and significance determined by bootstraps (1000 bootstraps for the SNP variants, and 100 bootstraps for the 491 832 sites due to computational limits) using modified functions in the R package diveRsity (Keenan et al. 2013). To examine the effects of laboratory-rearing on genetic diversity, we plotted the distributions of average heterozygosity in R (R Core Team 2014).

To investigate population differentiation, a global estimate of Weir and Cockerham’s (1984) F_ST_ with 99% bootstrap confidence intervals (10 000 bootstraps) was calculated in the R package diveRsity (Keenan et al. 2013). Pairwise F_ST_ values (Weir and Cockerham 1984) were calculated and significance determined using exact G tests implemented in genepop v4.3 (Rousset 2008) after Bonferroni correction for multiple comparisons (Dunn 1961). To test for genetic isolation by distance (Wright 1943), we performed a Mantel test (Mantel 1967) using 10 000 permutations on the regression of Slatkin’s (1995) linearized F_ST_ transformation (F_ST_/(1-F_ST_) onto the natural log of geographic distance (Rousset 1997) using the R package ade4 (Dray and Dufour 2007). Geographic distances were calculated using the geographic distance matrix generator (Ersts 2007).

We used the 1285 SNP variants to analyse population structure using a Bayesian clustering method in the program structure v2.3.4 (Pritchard et al. 2000). Variant data were converted from vcf to structure file format using pgdspider v2.0.8.2 (Lischer and Excoffier 2012). structure analysis was used to infer the number of genotypic clusters and assign individuals to clusters. Analyses were performed for all individuals (n=72) and separately for *P. xylostella* individuals only (n=69). For each analysis, we assumed the admixture model with correlated allele frequencies and the locprior model, specifying nine geographic populations. For each analysis, we performed ten independent structure runs for each value of K=1-10, where *K* is the number of genotypic clusters. For all runs, we used 500 000 burnins and 500 000 MCMC replicates. The optimal *K* was determined using the delta *K* method of Evanno et al. (2005) visualized in the program structure harvester (Earl and vonHoldt 2012). Individual and population Q-matrices (containing posterior probability of assignment to genotypic clusters) across replicate structure runs were aligned in the program clumpp v1.1.2 (Jakobsson and Rosenberg 2007) and visualized in distruct v1.1 (Rosenberg 2004).

### PCR genotyping assays

To examine the frequency of mutations associated with pyrethroid resistance, we performed PCR based genotyping assays for three point-mutations in the voltage gated sodium channel, T929I (*skdrl*), L1014F (*kdr*) and F1020S (*cdr*) according to Endersby et al. (2011). MyTaq polymerase (Bioline) was used for amplification in a Verity thermocycler (ABI).

To distinguish between *P. xylostella* and *P. australiana* lineages, we developed a PCR-RFLP genotyping assay using COI sequence published by Landry and Hebert (2013). Genomic DNA was amplified using a modified LCO1492_Px primer (5’-TCAACAAATCATAAAGATATTGG-3’) and HCO2198 (5’-TAAACTTCAGGGTGACCAAAAAATCA-3 ‘) (Folmer et al. 1994). Ten microliter reactions were run with 2 μL of MyTaq 10x buffer, 0.4 μL of each primer (10 μM stocks), 1 μL of DNA (approx. 5 ng) and 0.05 μL of MyTaq polymerase (Bioline). Samples were amplified at 95 °C for 2 minutes, then 35 cycles at 95°C for 10 seconds, 52°C for 20 seconds, 72°C for 30 seconds) followed by a 5 minute final extension at 72°C. PCR products were then digested at 37°C for 1 hour with *AccI* restriction enzyme with 2 μL Cutsmart Buffer and 1 unit of *AccI* (NEB) to a final volume of 20 μL. Following digestion, products were separated using agarose gel electrophoresis (1.5%). *P. xylostella* products are approximately 517 bp and 191 bp and *P. australiana* products are 348 bp and 360 bp (Figure 1C).

## RESULTS

### Population genetic analysis

Using RAD sequencing we identified a set of 491 832 variant and invariant sites, representing 0.146% of the *P. xylostella* reference genome, and a subset of 1285 SNP variants, for population genetic analysis. Individuals from nine field and laboratory-reared populations from Australia were genotyped at an average read depth of 45 (Table 1).

The 491 832 confident sites were used to generate a neighbour-joining phylogeny for all 72 individuals (Figure 1A,B). Seven individuals collected from South Australian locations, three from Calca, three from Nundroo and one from Picnic Beach, were confidently resolved with 100% consensus support. Three individuals, Calca-4,-6 and -7, showed particularly long branch lengths, although the branch length estimate for Calca-4 is inflated due to missing data (48% sites genotyped). We performed PCR mitochondrial genotyping assays to assess whether the dataset contained multiple *Plutella* lineages. The assays identified the three divergent individuals, Calca-4,-6 and -7, as a cryptic species, *P. australiana*, and confirmed that all other individuals (n=69) were *P. xylostella* (Figure 1C). These results based on RAD-Seq markers indicate strong divergence in the nuclear genome between the two Australian *Plutella* lineages, consistent with high mitochondrial sequence divergence already reported (Landry and Hebert 2013). Four basal *P. xylostella* individuals, Picnic Beach-5, -6, -9 and Nundroo-8, were also confidently resolved (100% consensus support), and relative branch length estimates suggest strong divergence from other *P. xylostella*. All remaining *P. xylostella* individuals could not be confidently resolved (<50% consensus support), despite some evidence of individuals grouping according to geographic location.

The levels of heterozygosity were used to assess genetic diversity within and among populations. For all genotyped sites, the average observed heterozygosity ranged from 0.0085 to 0.0150 for *P. xylostella* populations but was notably higher at 0.0220 for the three *P. australiana* individuals (Table 2). The average number of private alleles among *P. xylostella* populations ranged from 20-91, while three *P. australiana* individuals had a much higher average of 1476 private alleles (range 606-2063), consistent with strong divergence. For the variant SNP dataset, the average observed heterozygosity ranged from 0.3052 to 0.5115 for *P. xylostella* populations, and again was higher at 0.5619 for the three *P. australiana* individuals (Table 2). Private alleles were filtered out of the SNP variant dataset. Average levels of heterozygosity were variable within and among populations (Figure 2). Laboratory-reared populations, Esperance, Glenore Grove and Mt Sylvia, show the lowest levels of genetic diversity (Figure 2), and were founded with relatively low numbers of individuals, ranging from 25–59 (Table 1). The values for the inbreeding coefficient, FIS, were significantly different from zero for these three populations (Table 2).

The global estimate of FST across all *P. xylostella* populations and loci was significantly different from zero (F_ST_ = 0.0487, 99% CL 0.0187-0.0905), indicating significant genetic differentiation among the nine populations. The pairwise FST values indicate that the differentiation is associated with populations from the three regions, Western Australia, South Australia and Queensland (Table 3). However, patterns of differentiation did not clearly relate to geographic proximity and may reflect inbreeding in laboratory populations. Glenore Grove, for example, was highly differentiated from all other populations including those from Tenthill and Mt Sylvia in close proximity (18-28 km). Lower F_ST_ values among some South Australian populations suggests high levels of gene flow within this region, however these populations were also not differentiated from Tenthill in Queensland despite large geographic separation (1515-1819 km). The Mantel test for all pairwise population comparisons indicated a weak but significant effect of isolation by distance (n=69 individuals, r = 0.3048, p=0.0215) (Figure 2B). When analysed separately however, there was no relationship between genetic and geographic distance for pairwise comparisons of either ‘cage’ populations (n=33, F_2_-F_6_; Mantel’s r= 0.07388, p=0.36226) or ‘field’ populations (n=3, F_0_-F_1_; Mantel’s r= 0.88862, p=0.33307).

**Figure 1:**
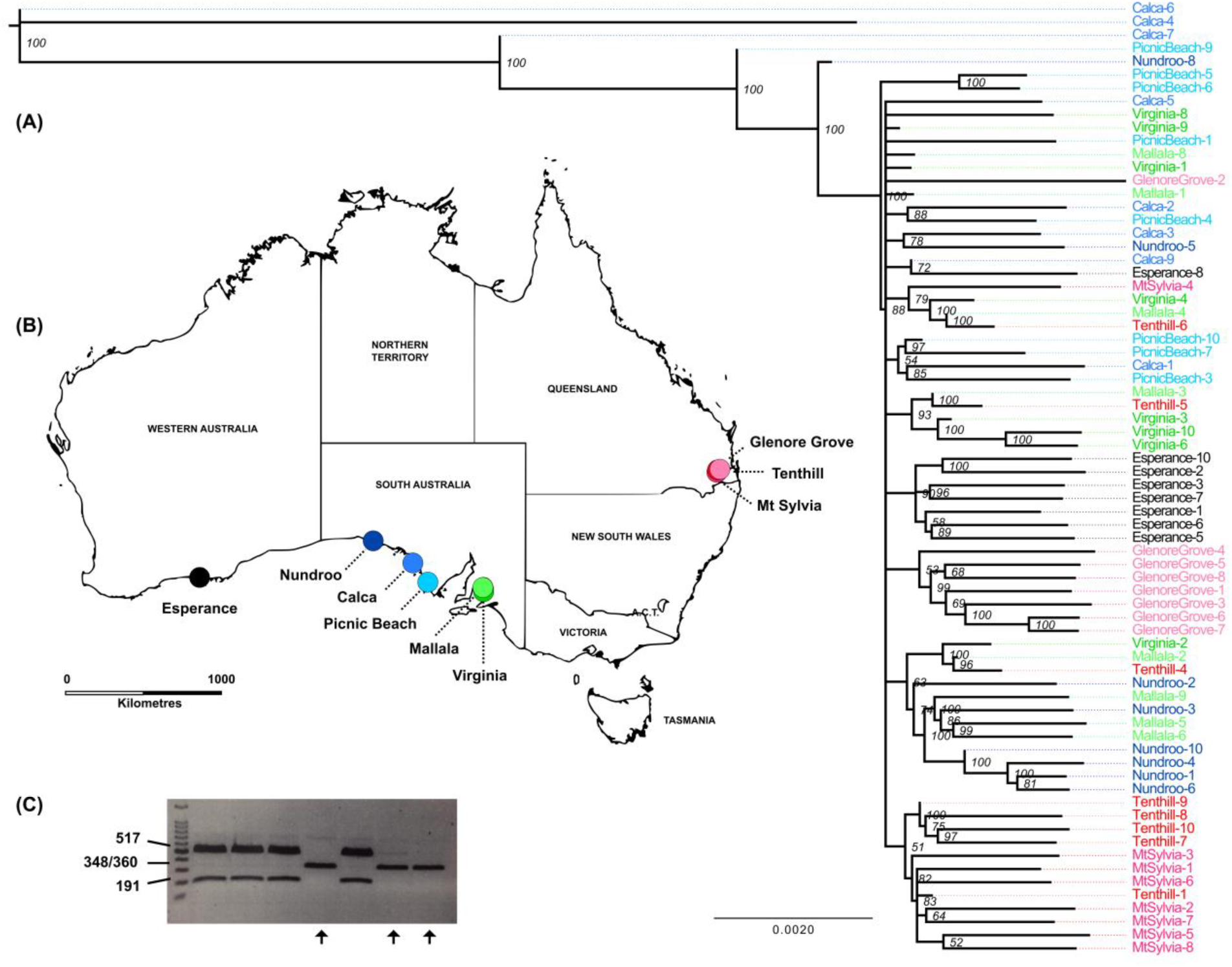
(A) Neighbour-joining consensus tree generated from 491 832 variant and invariant sites for 72 *Plutella* individuals, displaying nodes with at least 50% consensus support. Node labels are the percentage consensus support for 1000 bootstrap replicates. (B) Map of collection sites for nine *Plutella* populations from Western Australia (n=1), South Australia (n=5) and Queensland (n=3). (C) Gel electrophoresis image showing the results of a mitochondrial CO1 gene assay to distinguish the two *Plutella* lineages. *P. australiana* is genotyped as a single band, shown by black arrows, and *P. xylostella* as two bands.

**Figure 2:**
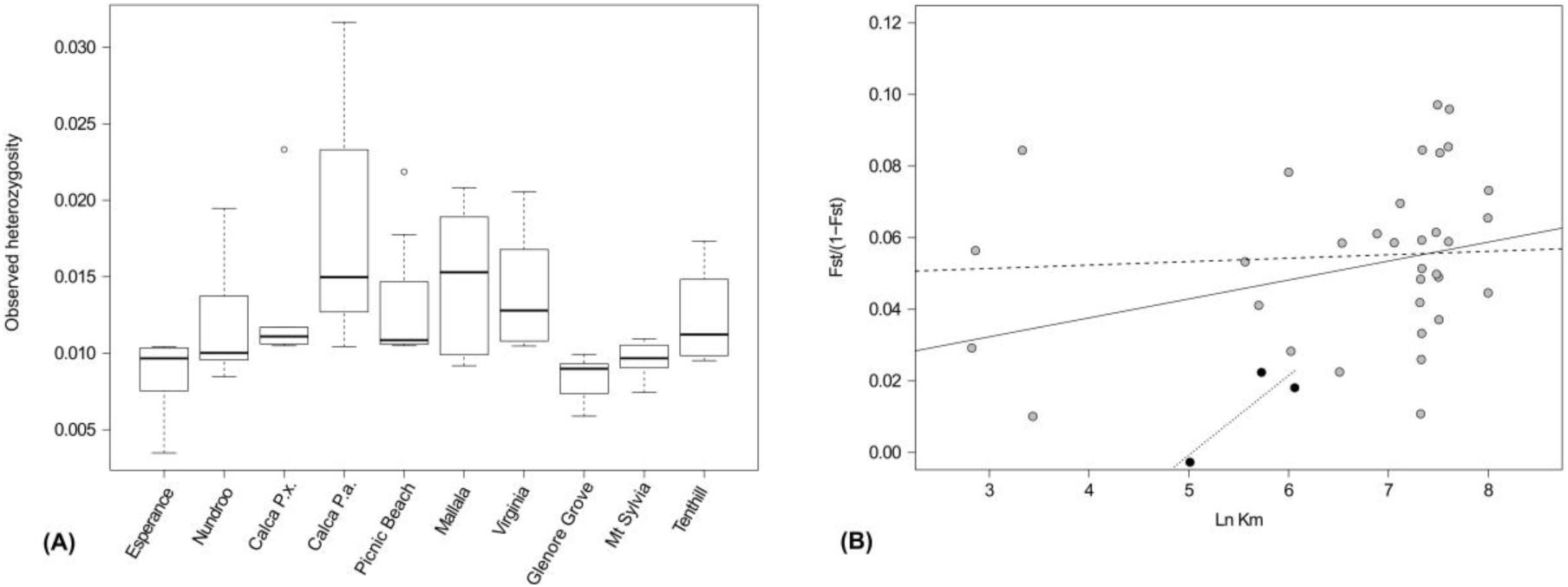
(A) Distributions of observed heterozygosity for *Plutella* populations. Each population contains eight individuals. Calca is split into two lineages, Calca P.x. (n=5) and Calca P.a. (n=3). (B) Regressions of Slatkin’s linearized genetic distance (F_ST_/1-F_ST_) against the natural log of geographic distance (Ln km) for all pairwise population comparisons of *P. xylostella* (n=69 individuals). Lines represent the fitted linear regression model for pairwise comparisons for ‘all populations’ (grey and black circles, solid line, y= 0.00529x+0.01646, Mantel’s r=0.3048, p=0.0215), ‘cage’ populations (grey circles, dashed line, y=0.00096x + 0.04854, Mantel’s r=0.07388, p=0.36226) and ‘field’ populations (black circles, dotted line, y=0.02224x-0.11197, Mantel’s r=0.88862, p=0.33307).

**Table 2:** Population statistics calculated for the SNP variant sites (top) and for all confidently called (GQ>30) variant and invariant sites (bottom) for populations of *Plutella* species collected from Australia. Populations each contain eight sequenced individuals with Calca split into *P. xylostella* (n=5) and *P. australiana* (n=3) individuals. Statistics include the population means for the number of individuals genotyped per locus (N), number of sites genotyped, site depth, number of sites unique to each population (private alleles), the proportion of observed (H_O_) and expected (H_E_) heterozygosity, and Wright’s inbreeding coefficient (F_IS_).

| Population | N | Sites<br>genotyped | Site depth | Private<br>alleles | $H_O$ | $H_E$ | $F_{IS}$ |
| --- | --- | --- | --- | --- | --- | --- | --- |
| **SNP variants (n=1 285)** |  |  |  |  |  |  |  |
| Esperance | 6.7 | 1073 | 32 | 0 | 0.3485 | 0.3650 | 0.0451 |
| Nundroo | 7.6 | 1215 | 54 | 0 | 0.4008 | 0.3763 | -0.0653 |
| Calca_Px | 4.8 | 1246 | 65 | 0 | 0.4327 | 0.3844 | -0.1256 |
| Calca_Pa | 2.0 | 855 | 34 | 0 | 0.5249 | 0.3233 | -0.6173 |
| Picnic Beach | 7.7 | 1238 | 73 | 0 | 0.3977 | 0.3821 | -0.0408 |
| Mallala | 7.7 | 1229 | 42 | 0 | 0.5619 | 0.4263 | -0.3181 |
| Virginia | 7.4 | 1194 | 30 | 0 | 0.5115 | 0.4032 | -0.2685 |
| Glenore Grove | 7.2 | 1158 | 34 | 0 | 0.3052 | 0.3469 | 0.1202 |
| Mt Sylvia | 7.3 | 1169 | 33 | 0 | 0.3490 | 0.3667 | 0.0482 |
| Tenthill | 7.3 | 1175 | 32 | 0 | 0.4827 | 0.4026 | -0.1991 |
| **All variant and invariant sites (n=491 832)** |  |  |  |  |  |  |  |
| Esperance | 7.5 | 462076 | 33 | 41 | 0.0090 | 0.0108 | **0.1715\*** |
| Nundroo | 7.8 | 479764 | 54 | 32 | 0.0120 | 0.0120 | 0.0041 |
| Calca_Px | 4.9 | 482269 | 65 | 91 | 0.0135 | 0.0129 | -0.0492 |
| Calca_Pa | 2.4 | 398538 | 39 | 1476 | 0.0220 | 0.0185 | -0.1923 |
| Picnic Beach | 7.9 | 482882 | 72 | 77 | 0.0132 | 0.0133 | 0.0113 |
| Mallala | 7.9 | 484254 | 43 | 20 | 0.0150 | 0.0129 | -0.1651 |
| Virginia | 7.8 | 480662 | 31 | 53 | 0.0142 | 0.0129 | -0.1017 |
| Glenore Grove | 7.6 | 464262 | 35 | 28 | 0.0085 | 0.0103 | **0.1800\*** |
| Mt Sylvia | 7.7 | 473837 | 34 | 30 | 0.0098 | 0.0112 | **0.1313\*** |
| Tenthill | 7.8 | 479215 | 33 | 20 | 0.0126 | 0.0118 | -0.0658 |

\* $F_{IS}$ values in bold are significantly different from zero according to bootstrapped 95% CL.

**Table 3:** Pairwise comparisons of genetic distance measured by Weir and Cockerham’s (1984) Fst (lower diagonal) and geographic distance in km (upper diagonal) for all population pairs of *P. xylostella*.

| | Esperance<br>($F_6$) | Nundroo<br>($F_2$) | Calca_Px<br>($F_0$) | Picnic<br>Beach ($F_1$) | Mallala<br>($F_4$) | Virginia<br>($F_0$) | Glenore<br>Grove ($F_5$) | Mt Sylvia<br>($F_5$) | Tenthill<br>($F_5$) |
| --- | --- | --- | --- | --- | --- | --- | --- | --- | --- |
| Esperance, WA |  | 979 | 1162 | 1234 | 1530 | 1534 | 2993 | 2968 | 2975 |
| Nundroo, SA | **0.0576\*\*\*** |  | 261 | 403 | 672 | 689 | 2021 | 1997 | 2004 |
| Calca_Px, SA | 0.0554 | 0.0506 |  | 150 | 413 | 429 | 1836 | 1811 | 1819 |
| Picnic Beach, SA | **0.0651\*\*\*** | **0.0726\*\*\*** | -0.0027 |  | 299 | 307 | 1793 | 1767 | 1776 |
| Mallala, SA | 0.0322 | 0.0220 | 0.0275 | 0.0395 |  | 31 | 1531 | 1504 | 1515 |
| Virginia, SA | 0.0489 | 0.0553 | 0.0178 | 0.0219 | 0.0100 |  | 1541 | 1514 | 1525 |
| Glenore Grove, Qld | **0.0682\*\*\*** | **0.0875\*\*\*** | **0.0772\*\*\*** | **0.0885\*\*\*** | **0.0560\*\*** | **0.0779\*\*\*** |  | 28 | 18 |
| Mt Sylvia, Qld | **0.0615\*\*\*** | **0.0787\*\*\*** | 0.0467 | **0.0579\*\*\*** | 0.0402 | 0.0462 | **0.0778\*\*\*** |  | 17 |
| Tenthill, Qld | 0.0427 | 0.0556 | 0.0358 | 0.0474 | 0.0108 | 0.0253 | **0.0534\*\*** | 0.0284 |  |

Significance of exact G tests after Bonferonni correction: $P < 0.05$ \*, $P < 0.001$ \*\*, $P < 0.0001$ \*\*\*

**Figure 3:**
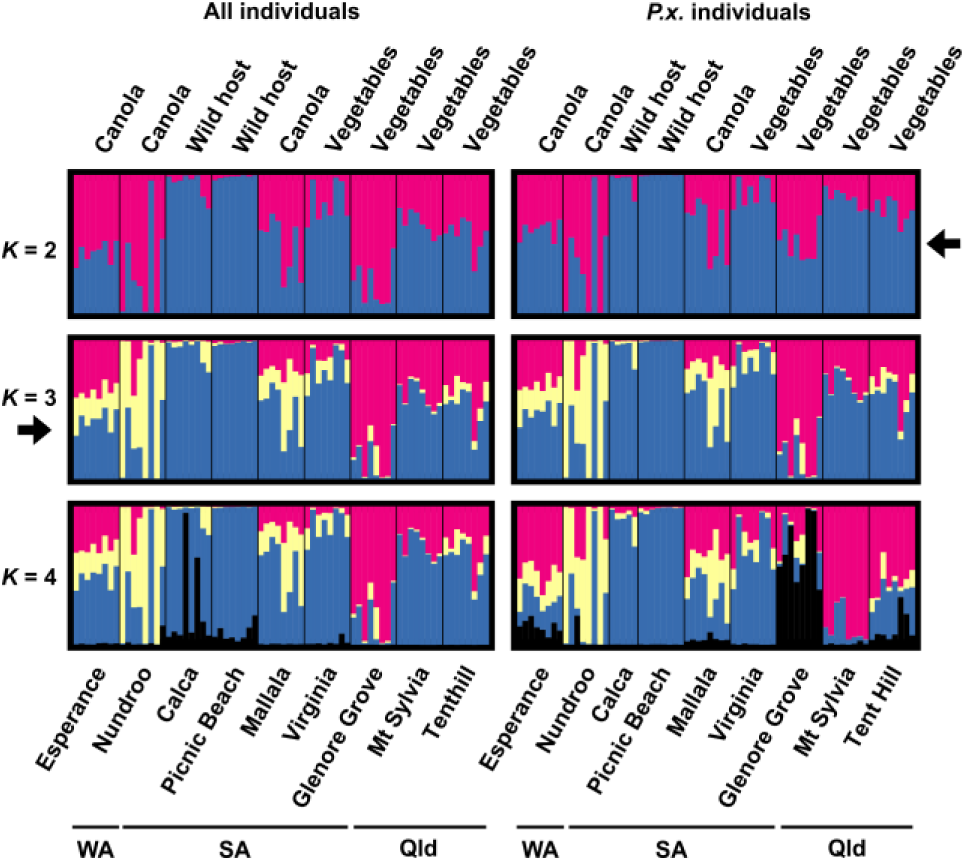
Posterior probability of assignment to inferred genotypic clusters, *K*, generated in the program structure for ‘all individuals’ (n=72), and separately for *‘P. xylostella* individuals’ (n=69) where three *P. australiana* individuals from the Calca population were excluded. The most likely *K* for ‘all individuals’ and ‘P.x. only’ is indicated by black arrows. Each vertical bar represents a single individual, populations are separated by black vertical lines, and different genotypic clusters for *K*=2-4 are represented by different colours.

We performed an analysis of population structure using a Bayesian clustering approach in the program structure. For the analysis of all population samples (n=72), the data most likely formed three genotypic clusters (*K*=3) (Figure 3). Inspection of structure barplots shows that individuals from Calca and Picnic Beach populations were assigned to similar genotypes clusters, however structure did not identify the three *P. australiana* individuals from Calca at that value of *K*. Although statistically less likely, *K* values of 2 and 4 are also presented. At *K*=4, the *P. australiana* individuals are more clearly resolved, as seen by tall black bars for individuals Calca-4 and -6 (Figure 3A). Moderate sharing of the black-coloured genotype cluster occurs across all individuals from Calca and Picnic Beach, and rarely elsewhere. For the analysis excluding the three *P. australiana* individuals, the most likely number of clusters was reduced to two (*K*=2). At this value of *K*, a high degree of admixture is evidence among most populations, as seen by sharing of pink and blue-coloured genotypic clusters. The two populations collected from wild hosts in a similar region, Calca and Picnic Beach, share similar genotypic clusters. At *K*=3, the assignment of individuals to genotypic clusters is consistent with patterns of population differentiation inferred from pairwise F_ST_ values (Table 3).

### Frequency of insecticide resistance alleles

We examined the frequency of mutations associated with pyrethroid resistance among nine populations collected from canola crops, *Brassica* vegetable crops and wild hosts. The average frequencies for *skdrl, kdr* and *cdr* were 0.29, 0.51 and 0.27 respectively (Appendix). The frequencies were comparable among populations for *skdrl* (range 0.2–0.38) and *kdr* (range 0.25–0.83), but more variable for *cdr* (0.05–0.6). Interestingly, the populations collected from wild hosts, Calca and Picnic Beach, had a moderately high frequency of the *cdr* mutation relative to most other populations. In contrast, the three *P. australiana* samples were all found to be susceptible for *skdrl* and *cdr* however the *kdr* assay failed, possibly due to variation in primer binding sites.

## DISCUSSION

RAD sequencing was used to identify thousands of SNP markers and hundreds of thousands of invariant loci from across the genome of two *Plutella* species. These markers facilitated an initial assessment of genetic diversity within and among nine *Plutella* populations collected from different locations and host plants across Australia.

Analysis of RAD-Seq markers and mitochondrial genotyping identified three individuals of a cryptic *Plutella* lineage, *P. australiana*, among our 72 *Plutella* individuals. The relative branch length estimates in the neighbouring-joining tree, differences in heterozygosity and high numbers of private alleles within the *P. australiana* populations (albeit only three individuals) provide the first evidence that the *P. australiana* and *P. xylostella* lineages are strongly divergent in their nuclear genomes. Among the three *P. australiana* individuals, it was interesting to note that individual Calca-7 had the fewest number of private alleles (606, compared to 1760 and 2062) despite having the highest heterozygosity (3.2%, compared to 1.5% and 1.04%). The phylogeny also shows this individual most closely related to *P. xylostella* individuals Picnic Beach-9 and Nundroo-8, which grouped separately from all other *P. xylostella* individuals. As yet, the potential for hybridization between these lineages remains to be tested.

The original discovery of *P. australiana* in Australia was made through sequencing the mitochondrial COI gene from moths collected in light traps, rather than from known host plants (Landry and Hebert 2013). Hence, the fundamental biology of this species and its potential pest status remain to be understood. In our study, individuals of *P. australiana* and *P. xylostella* were collected from a wild Brassicaceous host, *Diplotaxis* sp., at the same location and date, indicating that these lineages can coexist in similar environments and exploit at least one common host. Considering that there have been several previous genetic studies of *P. xylostella* in Australia (Endersby et al. 2006, Pichon et al. 2006, Roux et al. 2007, Endersby et al. 2008), including mitochondrial markers (Saw et al. 2006), the discovery of this novel lineage only recently is intriguing. It is possible that differences in sampling strategies (direct sampling from plants vs trapping), times or locations between studies, or differences in the biology of *Plutella* lineages (e.g. host range), meant that *P. australiana* was not collected in previous studies. Alternatively, some molecular markers designed for *P. xylostella* may not amplify efficiently in *P. australiana*. These questions require further investigation.

We examined genetic diversity in field and laboratory-reared populations of *P. xylostella*. Significantly reduced heterozygosity was observed in the laboratory populations, Glenore Grove, Mt Sylvia and Esperance, as measured by the inbreeding co-efficient, FIS (Table 2). These populations were founded by 25-59 individuals at the F1 generation and then reared for five to six generations. The population from Mallala was also reared for five generations but established from a higher number of individuals (n=173), and maintained higher levels of heterozygosity in culture, comparable with the field (F0) populations.

We assessed genetic structure among our population samples using a range of approaches. The global estimate of F_ST_ (0.0487, 99% CL 0.0187-0.0905) indicated significant genetic differentiation among our populations. Pairwise F_ST_ comparisons showed that most differentiation was associated with the three most inbred laboratory populations, Glenore Grove, Mt Sylvia, Esperance, but also Nundroo (F_2_). There was no evidence for isolation by distance in pairwise population comparisons among laboratory populations only. Hence, the estimates of genetic isolation are inflated by inbreeding in population cages. The structure analysis for all population samples (n=72) inferred that individuals most likely form three genotypic clusters, however failed to resolve the two lineages at that value of *K*. This result could reflect that alleles unique to *P. australiana* individuals were filtered out of the variant SNP dataset used for this analysis. Removing *P. australiana* reduced the optimal value of *K* to two. At this *K* value, a large degree of admixture was observed among populations, supporting the neighbor-joining phylogeny which failed to clearly resolve clusters for the majority of *P. xylostella* individuals (<50% consensus support). Overall, these findings are consistent with high levels of gene flow previously reported for Australian populations of diamondback moth (Endersby et al. 2006).

The frequency of pyrethroid resistance alleles has previously been documented for Australian *P. xylostella* populations collected from 2003–2005 (Endersby et al. 2011). Endersby et al. (2011) reported average resistance allele frequencies for *skdrl* (0.139), *kdr* (0.609) and *cdr* (0.305), however considerable spatial variation was observed. To assess the potential change in frequencies over time, we re-examined these frequencies from populations collected between 2012 and 2014 (Table 1, 4). The average frequency for *skdrl, kdr* and *cdr* were 0.29, 0.51 and 0.27 respectively, which is comparable between studies for *kdr* and *cdr*, however somewhat higher for *skdrl*. The stability of these frequencies may reflect that synthetic pyrethroid insecticides continue to be widely used to control a range of invertebrate pests in Australian crops.

## CONCLUSION

RAD-Seq is a powerful method for generating SNP markers for population genetic studies in the diamondback moth. To aid biological inference, we recommend that future studies focus on field sampling design and wherever possible strive to use field-collected (F_0_) populations to adequately represent genetic diversity.

## Acknowledgements

We thank Kevin Powis, Fred Bartholomaeus and Greg Baker (South Australian Research & Development Institute) for population samples and meta-data, Claudia Junge for helpful advice on the analysis, and Eliska Zlamalova for assistance with PCR genotyping. KDP is funded by The University of Adelaide and the Grains Research & Development Corporation (UA00146). SWB is funded by the Australian Research Council (DP120100047, FT140101303).

## Appendix

**Table 4:** Frequency of resistance alleles for three known point mutations in the voltage-gated sodium channel (Endersby et al. 2011) in nine populations of *P. xylostella:* super-kdr-like (*skdrl*), knockdown resistance (*kdr*), and crashdown (*cdr*).

| Locality | Host type | Individuals genotyped (*skdrl*/*kdr*/*cdr*) | Resistance allele frequency |  |  |
| --- | --- | --- | --- | --- | --- |
|  |  |  | *skdrl* | *kdr* | *cdr* |
| Esperance, Western Australia | Canola | 7/6/8 | 0.36 | 0.58 | 0.06 |
| Nundroo, South Australia | Canola | 10/10/10 | 0.20 | 0.25 | 0.1 |
| Calca, South Australia | Wild host | 9/9/9 | 0.38 | 0.63 | 0.31 |
| Picnic Beach, South Australia | Wild host | 12/9/12 | 0.33 | 0.72 | 0.29 |
| Mallala, South Australia | Canola | 10/6/10 | 0.35 | 0.67 | 0.05 |
| Virginia, South Australia | Vegetables | 10/10/10 | 0.30 | 0.40 | 0.60 |
| Glenore Grove, Queensland | Vegetables | 8/5/8 | 0.25 | 0.60 | 0.19 |
| Mt Sylvia, Queensland | Vegetables | 8/6/9 | 0.31 | 0.83 | 0.06 |
| Tenthill, Queensland | Vegetables | 10/10/10 | 0.30 | 0.40 | 0.20 |
| **By Host type:** |  |  |  |  |  |
| Canola (n. pops = 3) | C | 27/22/28 | 0.30 | 0.45 | 0.07 |
| Wild hosts (n. pops = 2) | W | 21/18/21 | 0.35 | 0.65 | 0.30 |
| Vegetables (n. pops = 4) | V | 36/31/37 | 0.29 | 0.51 | 0.27 |

